# Effect of tag attachment on the flight performance of five raptor species

**DOI:** 10.1101/2021.09.15.460505

**Authors:** Arianna Longarini, Olivier Duriez, Emily Shepard, Kamran Safi, Martin Wikelski, Martina Scacco

## Abstract

Bio-logging devices play a fundamental and indispensable role in movement ecology studies, particularly in the wild. However, researchers are becoming increasingly aware of the influence that attaching devices can have on animals, particularly on their behaviour, energy expenditure and survival. The way a device is attached to an animal’s body has also potential consequences for the collected data, and quantifying the type and magnitude of such potential effects is fundamental to enable researchers to combine and compare data from different studies, as much as it is to improve animal welfare.

For over two decades, large terrestrial birds have been in the focus of long-term movement ecology research, employing bio-logging devices attached with different types of harnesses. However, comparative studies investigating the effects of different harness types used on these species are scarce.

In this study, we tested for potential differences in data collected by two commonly used harness types, backpack and leg-loop, on the flight performance of 10 individuals from five raptor species, equipped with high resolution bio-logging devices, in the same area and time. We explored the effect of harness type on vertical speed, horizontal speed, glide ratio, height above sea level, distance travelled, proportion of soaring and flapping behaviour, and VeDBA (a proxy for energy expenditure) between and within individuals, all used as fine-scale measures of flight performance. Birds equipped with leg-loops climbed up to 0.65 ms^−1^ faster, reached 19% greater heights while soaring, and spent less time in active flight compared to birds equipped with backpacks, suggesting that backpack harnesses, compared to leg-loops, might cause additional drag affecting the birds’ flight performance. A lower rate of sinking while gliding, a slightly higher glide ratio, higher horizontal speed while soaring, and lower VeDBA, were also indicative of less drag using leg-loops. Our results add to the existing literature highlighting the design-related advantages of leg-loops, and support the use of leg-loops as a better alternative to backpack harnesses for large soaring birds, when possible. Our study also highlights how apparently small changes in device attachment can lead to notable improvements in tagging practice, with implications for animal welfare, data interpretation and comparability.

## Background

The recent advances in the movement ecology field are sparked by the growing possibilities to remotely measure the movement and behaviour of animals in the wild. The use of bio-logging devices, such as GPS loggers, accelerometers and internal sensors, allow us to record an unprecedented amount of quantitative information concerning the movement and behaviour of an animal, its physiological condition and its environmental context (Williams et al. 2020).

Despite the fundamental role of bio-logging techniques in movement ecology studies, and the ensuing gain in knowledge, researchers are increasingly aware of the potential effects that bio-logging devices can have on animal behaviour and survival. Flying animals are in that respect of special concern. Bio-logging is fundamental to studying their long-distance movements; however, the added weight of a device can challenge their ability to remain aloft. In addition, the device’s shape and position can increase drag during flight, and its attachment, when done without the necessary diligence, create discomfort around the wings. Recent studies suggested an adverse effect of bio-logging on several aspects of avian behaviour and ecology (Barron et al. 2010), including lower recapture and survival rate, a decreased likelihood of nesting success, nesting productivity and nesting propensity, changes in foraging trip duration, as well as an increase in energy expenditure, predation risk and death (Calvo & Furness 1992, Culik et al. 1993, Ballard et al. 2001, Zuberogoitia et al. 2012, Trefry et al. 2013, Vandenabeele et al. 2014, Watanuki et al. 1992, Miller & Davis 1993, Navarro et al. 2008, Taylor et al. 2010). However, other studies did not find neither short- nor long-term differences in reproductive success, survival, activity budget and return rate at the colonies, attributable to the attachment of bio-logging devices Hamel et al. (2004), Thaxter et al. (2016). Very few studies investigated the effect of the use of bio-logging on flight performance, but some highlighted how birds can reach different flight speeds depending on tag placement (Gessaman & Nagy 1988, Wilson & Culik 1994, Curk, T., Scacco, M. et al. 2021). More commonly, studies focus on the effects of device weight relative to the animal’s body mass, the device shape and induced drag depending on the medium in which the animal moves (Vandenabeele et al. 2012, Wilson & Culik 1994, Bowlin et al. 2010), the device position relative to the centre of mass (Wanless et al. 1989, Powell et al. 1998, Vandenabeele et al. 2014) and the material of the harness used to attach the device (Barron et al. 2010, Vandenabeele et al. 2013). Harnesses are indispensable for long-term bio-logging studies (Naef-Daenzer 2007). Large terrestrial birds (raptors and large soaring birds), including many endangered species, are often the subject of such important research, but few studies investigated the effect of harness type on these species. In fact, studies investigating the effect of the type of harness used to attach a device are mostly concentrated on waterbirds like penguins, waterfowl and seabirds. In addition, studies comparing different types of harnesses on individual species are overall scarce (Thaxter et al. 2014), especially in the case of terrestrial birds, and are usually based on few individuals (but see (Steenhof et al. 2006)).

Long-term studies on raptors usually employ backpack-type (thoracic) harnesses (Thaxter et al. 2014, Naef-Daenzer 2007, Anderson et al. 2020). Some studies on raptors found that this type of harness causes irritation under the wings, physical discomfort and increases preening behaviour (Booms et al. 2011, Stahlecker et al. 2015, Anderka & Angehrn 1992). Other studies showed that backpack harnesses decreased the survival in Spotted Owls *Strix occidentalis* (Paton et al. 1991) and Prairie Falcons *Falco mexicanus* (Steenhof et al. 2006). Birds equipped with this harness type are also at risk of entangling their wings, especially if the harness is too loose. On the contrary, if too tight, this might inhibit the action of flight muscles or the deposition of fat (Naef-Daenzer 2007, Thaxter et al. 2014). In addition, the design of backpack harnesses, consisting of two loops connected over the sternum, makes it difficult to impossible for the harness to fall off, in case of rupture of one of the loops. This will force the bird to unnecessarily keep carrying a damaged harness, in an improper position and often failing to work, yet hindering the bird’s movements. Backpack harnesses are still widely used, particularly on terrestrial birds, and continue to provide indispensable insight into the movement of animals and their interactions with the environment, offering the basis for effective conservation and mitigation measures. However, alternative harness types deserve some attention. In recent years, leg-loop harnesses (or Rappole-type harnesses), originally introduced for passerines, have started being used on larger species too, especially seabirds (Thaxter et al. 2014, Rappole & Tipton 1991). Leg-loop harnesses consist of two loops, each passing around the bird’s thighs, with the device resting on its lower back. Their design leaves wings, flight muscles and major fat deposits untouched. It also reduces the risk of entanglement, and contrary to backpacks, if one side of the harness gets damaged, a leg-loop harness will fall off. Leg-loops, albeit certainly also representing a burden on the studied individuals, might therefore be considered a valid alternative to backpack harnesses. However, the applicability of leg-loops is not universal, as for species with short thighs it isn’t a safe attachment method (Rappole & Tipton 1991, Naef-Daenzer 2007). Also, due to the position on the lower back, one study reported difficulties in solar-charging the battery of devices attached with leg-loop design (Thaxter et al. 2014). Therefore as for backpack harnesses, the applicability of leg-loops has to consider the morphological, demographic, and behavioural specifics of the species studied, with the goal of minimising impact on the natural behaviour of the individuals as an ethical responsibility, while also maximizing data quality and acquisition.

Leg-loop harnesses have been recently used on raptor species, but to our knowledge no study investigated their long-term reliability compared to the more commonly employed backpack harnesses, nor their short-term effects on the birds’ behaviour and flight performance. In this respect, despite the advantages of leg-loop harnesses, their design forces the device in a position that, compared to backpack harnesses, is further away from the bird’s centre of mass, and could cause higher energetic costs Vandenabeele et al. (2014). This potential consequence has hitherto been neglected and would be important to investigate.

In this study, we tested the effects of backpack and leg-loop harnesses on the flight performance of 10 individuals from five raptor species, equipped with high resolution bio-logging devices. Specifically, we explored the effect of using backpack *vs* leg-loop attachment on vertical speed, horizontal speed, glide ratio, height above sea level, distance travelled, proportion of soaring and flapping behaviour, and VeDBA (Vectorial Dynamic Body Acceleration, a proxy for energy expenditure (Wilson et al. 2020)), all used as measures of flight performance. The species involved were: grif-fon vulture (*Gyps fulvus*), Rüppell’s vulture (*Gyps rueppelli*), Himalayan griffon vulture (*Gyps himalayensis*), tawny eagle (*Aquila rapax*) and black kite (*Milvus migrans*). These five species are characterised by different morphology, spanning a range of body masses from 0.8 to 8.4 Kg and wing spans from 1.38 to 2.8 m. The study was performed in a falconry park during a week of data collection, consisting of three flight sessions per day. During each flight session, we equipped the birds with high resolution GPS and accelerometry devices. The falconry park provided the unique setting of a common-garden experiment: all 10 individuals from the five species flew simultaneously in the same area, thus experiencing roughly the same environmental conditions; this minimized confounding factors related to the environmental context and facilitated comparisons across species. It also allowed us, during subsequent days, to collect data on the same individuals while attaching devices on them with one or the other harness type. This helped minimizing differences in flight performance related to the individuals’ behaviour rather than on the harness type. Moreover, all individuals were used to be handled on a daily basis, which likely reduced the stress usually associated with handling wild birds.

## Results

### Analysis of the behavioural segments

Between the 28^th^ of June and the 1^st^ of July 2018, in a falconry centre in Rocamadour (France), we collected GPS and tri-axial accelerometry (ACC) data on 10 individuals from five raptors species: Eurasian griffon vulture (n=4), Rüppell’s vulture (n=1), Himalayan griffon vulture (n=2), tawny eagle (n=2) and black kite (n=1). GPS and ACC devices were attached to the birds using harnesses fitted either as a leg-loop or as a backpack.

The unit of this analysis was the behavioural segment, classified based on the GPS data as either soaring or gliding, and based on the ACC data as either passive or active flight. Our data included a total of 2172 observations (37 for the control individual, 2135 for the treatment individuals), where each observation corresponded to the average flight parameters of one behavioural segment. The five flight parameters associated to each segment were: mean vertical speed, mean horizontal speed, glide ratio (as the horizontal distance covered per unit of vertical distance dropped), maximum height a.s.l. and mean VeDBA; their distribution relative to harness type, for both the control and treatment groups, is shown in figures 1 and 2.

**Figure 1:**
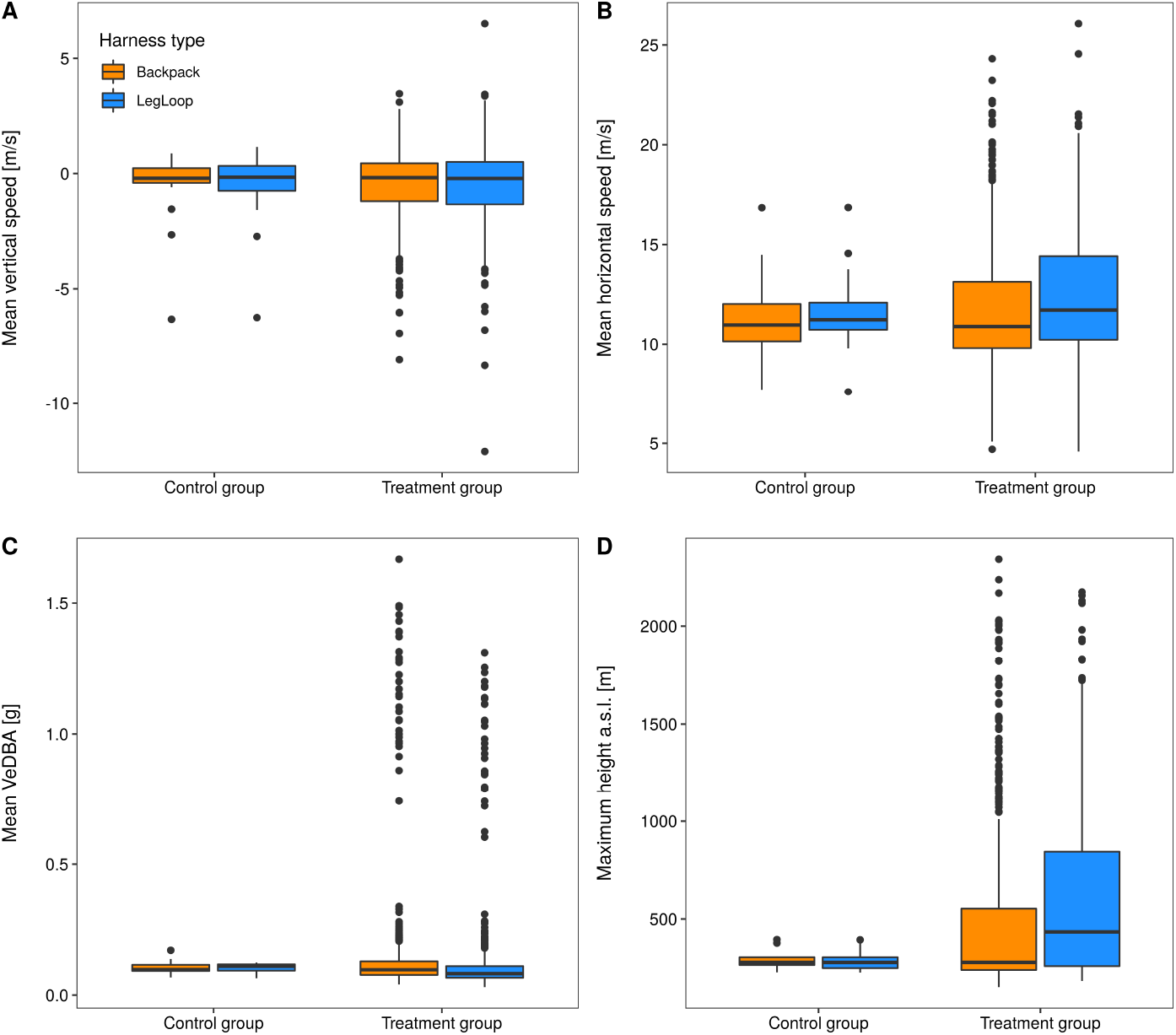
Average (A) vertical speed, (B) horizontal speed, (C) VeDBA, and (D) maximum height a.s.l. per behavioural segment, in the control bird and the treatment individuals. Different colours differentiate between individuals equipped with backpack and leg-loop harnesses.

**Figure 2:**
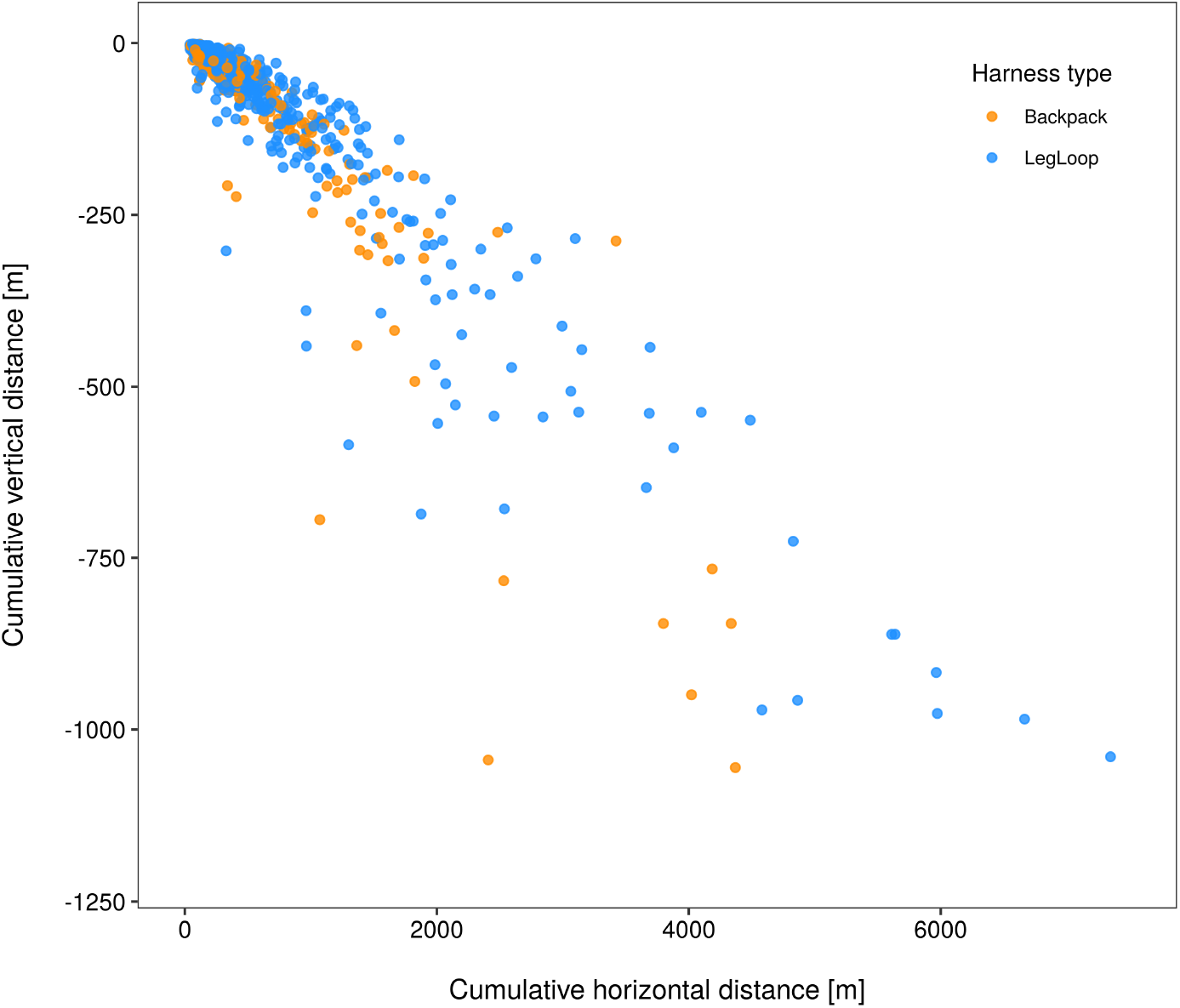
Cumulative distance covered in the horizontal plane relative to the cumulative vertical distance dropped per gliding segment. Different colours differentiate between individuals equipped with backpack and leg-loop harnesses.

### Control group

The dataset of the control individual included two flight sessions during which two devices were attached simultaneously to the bird, one with a leg-loop and one with a backpack harness, and collected a total of 37 observations (18 backpack and 19 leg-loop). All behavioural segments included in this dataset were classified as passive behaviour (either soaring or flapping). Using the two-sided Wilcoxon tests we detected no significant difference in the distribution of the five flight parameters between backpack and leg-loop segments, indicating that the accuracy of the information measured by the devices was not affected by their position [mean vertical speed: W = 170.5, *p* = 1; mean horizontal speed: W = 145, *p* = 0.44; glide ratio: W = 27, *p* = 0.75; maximum height a.s.l.: W = 180.5, *p* = 0.78; mean VeDBA: W = 146, *p* = 0.46].

### Treatment group

The dataset of the treatment group included 92 flight sessions from 10 individuals. During each flight session, individuals were equipped with either a leg-loop or a backpack harness. The complete dataset included a total of 2135 observations (789 backpack and 1346 leg-loop). The flight parameters mean vertical speed and mean horizontal speed were each used as response variable in two LMMs (one for the soaring and one for the gliding flight segments, including 1208 and 927 observations respectively). Maximum height a.s.l. was also analysed in two separate LMMs but the number of observations was halved (every second observation excluded) to reduce temporal auto-correlation, obtaining 604 soaring segments and 464 gliding segments. Glide ratio was analysed during gliding segments only (859 observations). VeDBA was only considered during passive flight, given the low number of active flight segments included in our dataset (N = 69); also in this case the dataset was halved to reduce temporal auto-correlation, obtaining 1037 observations. All models’ results listed below, unless otherwise specified, show estimate ± st.err.

In the vertical speed model associated to soaring, the effect of harness type differed between species, the interaction term being significant compared to the null model [*χ*^2^ = 15.17, *p* = 0.004]. All vultures species equipped with leg-loops reached significantly higher vertical speeds while soaring, up to 0.65 ms^−1^ higher (Rüppell’s vulture), compared to the backpack group [leg-loop:Griffon vulture = 0.51 ± 0.20; leg-loop:Himalayan vulture = 0.39 ± 0.21; leg-loop:Rüppell’s vulture = 0.65 ± 0.25], while the effect on the black kite and the tawny eagle was statistically non significant (Table 1). In the gliding model the effect of harness type did not differ between species [*χ*^2^ = 4.99, *p* = 0.29] but overall all species showed a significant increase in vertical speed (lower sinking rate) when equipped with leg-loops [leg-loop = 0.15 ± 0.08] (Table 1).

**Table 1:** Output of the LMM with mean vertical speed included as dependent variable, number of fixes, hour of the day, harness type and species as fixed terms, individual identity and date as random intercepts. The interaction term between harness type and species was non significant in the gliding model and therefore excluded.

|  | Soaring segments | Gliding segments |
| --- | --- | --- |
| **Fixed effects** |  |  |
| *Estimate (St. Err.)* |  |  |
| Intercept | 0.39 (0.15)* | -0.99 (0.33)* |
| Leg-loop | -0.29 (0.19) | 0.15 (0.08)* |
| Tawny eagle | -0.18 (0.19) | -0.71 (0.43) |
| Griffon vulture | -0.22 (0.15) | -0.40 (0.37) |
| Himalayan vulture | -0.20 (0.16) | -0.33 (0.40) |
| Rüppell's vulture | -0.50 (0.17)* | -0.85 (0.46) |
| Hour | -0.08 (0.02)*** | 0.02 (0.03) |
| Number of fixes | 0.004 (0.0002)*** | -0.007 (0.0005)*** |
| Leg-loop*Tawny eagle | 0.02 (0.26) |  |
| Leg-loop*Griffon vulture | 0.51 (0.20)* |  |
| Leg-loop*Himalayan vulture | 0.39 (0.21). |  |
| Leg-loop*Rüppell's vulture | 0.65 (0.25)** |  |
| **Random effects (N. groups)** |  |  |
| *Intercept St. Dev.* |  |  |
| Individuals | 0.05 (10) | 0.30 (10) |
| Date | 0.16 (7) | 0.15 (7) |
| Observations | 1208 | 926 |
| Marginal R<sup>2</sup> | 0.25 | 0.19 |
| Conditional R<sup>2</sup> | 0.29 | 0.28 |
| . p<0.1; *p<0.05; **p<0.01; ***p<0.001 |  |  |

In the case of the horizontal speed, in the soaring segments the effect of harness type did not differ between species [*χ*^2^ = 2.95, *p* = 0.57] but overall, all individuals showed a significant increase in horizontal speed when equipped with leg-loops and were predicted to fly up to 0.44 ms^−1^ faster when equipped with leg-loops [leg-loop = 0.08 ± 0.02] (Table 2). In the horizontal speed model associated to gliding, the different species showed a different response to harness type [*χ*^2^ = 10.38, *p* = 0.034], but this difference was statistically significant only in the Himalayan vulture and in the black kite. Himalayan vultures were predicted to glide 2.12 ms^−1^ slower when wearing a leg-loop [leg-loop:Himalayan vulture = −2.12 ± 0.80]. On the contrary, the smallest species, the black kite, showed a significant increase in horizontal speed when equipped with a leg-loop [leg-loop = 1.37 ± 0.70] (Table 2).

**Table 2:** Output of the LMM with the mean horizontal speed included as dependent variable, number of fixes, hour of the day, harness type and species as fixed terms, individual identity and date as random intercepts. In the soaring model the dependent variable was square root transformed. The interaction term between harness type and species was non significant in the soaring model and therefore excluded.

|  | Soaring segments | Gliding segments |
| --- | --- | --- |
| **Fixed effects** |  |  |
| *Estimate (St. Err.)* |  |  |
| Intercept | 2.63 (0.095)*** | 9.19 (1.13)** |
| Leg-loop | 0.08 (0.02)*** | 1.37 (0.70)* |
| Tawny eagle | 0.30 (0.11)* | 2.92 (1.68) |
| Griffon vulture | 0.76 (0.10)** | 4.90 (1.28)* |
| Himalayan vulture | 0.68 (0.11)** | 4.61 (1.37)* |
| Rüppell's vulture | 0.60 (0.12)** | 3.49 (1.57). |
| Hour | -0.007 (0.007) | -0.04 (0.07) |
| Number of fixes | 0.00009 (0.00008) | -0.009 (0.002)*** |
| Leg-loop*Tawny eagle |  | -2.3 (1.46) |
| Leg-loop*Griffon vulture |  | -1.16 (0.75) |
| Leg-loop*Himalayan vulture |  | -2.12 (0.80)** |
| Leg-loop*Rüppell's vulture |  | -0.66 (0.98) |
| **Random effects (N. groups)** |  |  |
| *Intercept St. Dev.* |  |  |
| Individuals | 0.08 (10) | 1.03 (10) |
| Date | 0.09 (7) | 0.49 (7) |
| Observations | 1208 | 927 |
| Marginal R<sup>2</sup> | 0.29 | 0.18 |
| Conditional R<sup>2</sup> | 0.38 | 0.30 |
| . p<0.1; *p<0.05; **p<0.01; ***p<0.001 |  |  |

In the glide ratio model the effect of harness type did not differ between species [*χ*^2^ = 2.52, *p* = 0.64] but overall, birds equipped with leg-loops showed a small and slightly significant increase in glide ratio [leg-loop = 0.16 ± 0.08]. This translates in about 1.07 m increase in horizontal distance covered per meter of drop for birds wearing leg-loops (Table 3).

**Table 3:** Output of the LMM with the square root of the glide ratio included as dependent variable, number of fixes, hour of the day, harness type and species as fixed terms, individual identity and date as random intercepts. The model included only gliding segment with vertical speed < 0.2 ms^−1^. The interaction term between harness type and species was not significant and therefore excluded.

|  | Gliding segments |
| --- | --- |
| **Fixed effects** |  |
| *Estimate (St. Err.)* |  |
| Intercept | 3.35 (0.20)*** |
| Leg-loop | 0.16 (0.08)* |
| Tawny eagle | -0.41 (0.28) |
| Griffon vulture | 0.09 (0.22) |
| Himalayan vulture | 0.003 (0.23) |
| Rüppell's vulture | -0.30 (0.26) |
| Hour | 0.009 (0.03) |
| Number of fixes | -0.005 (0.0005)*** |
| **Random effects (N. groups)** |  |
| *Intercept St. Dev.* |  |
| Individuals | 0.15 (10) |
| Date | 0.16 (7) |
| Observations | 859 |
| Marginal $R^2$ | 0.12 |
| Conditional $R^2$ | 0.16 |
| . $p < 0.1$; *$p < 0.05$; **$p < 0.01$; ***$p < 0.001$ | |

In both models predicting the maximum height a.s.l. the effect of harness type did not differ between species, the interaction terms being non significant compared to the null models [soaring: *χ*^2^ = 9.42, *p* = 0.05; gliding: *χ*^2^ = 6.86, *p* = 0.14]. Both models showed that birds reached higher altitudes when equipped with leg-loops. This effect was highly significant during soaring, associated to a 19% increase in altitude [soaring: leg-loop = 0.19 ± 0.05], and slightly significant during gliding [gliding: leg-loop = 0.11 ± 0.006] (Table 4).

**Table 4:** Output of the LMM with the log of the maximum height a.s.l. included as dependent variable, number of fixes, hour of the day, harness type and species as fixed terms, individual identity and date as random intercepts. Both models were run on a subset of the dataset (every second observation was discarded) to reduce temporal auto-correlation. In both models, the interaction term between harness type and species was non or slightly significant and therefore excluded.

|  | Soaring segments | Gliding segments |
| --- | --- | --- |
| **Fixed effects** |  |  |
| *Estimate (St. Err.)* |  |  |
| Intercept | 5.31 (0.13)*** | 5.45 (0.29)*** |
| Leg-loop | 0.19 (0.05)*** | 0.11 (0.059). |
| Tawny eagle | 0.24 (0.16) | 0.32 (0.36) |
| Griffon vulture | 0.55 (0.13)** | 0.62 (0.32) |
| Himalayan vulture | 0.47 (0.13)* | 0.46 (0.34) |
| Rüppell's vulture | 0.35 (0.15). | 0.65 (0.40) |
| Hour | -0.03 (0.02) | -0.01 (0.02) |
| Number of fixes | 0.003 (0.0002)*** | 0.004 (0.0004)*** |
| **Random effects (N. groups)** |  |  |
| *Intercept St. Dev.* |  |  |
| Individuals | 0.07 (10) | 0.27 (10) |
| Date | 0.16 (7) | 0.09 (7) |
| Observations | 604 | 464 |
| Marginal R<sup>2</sup> | 0.27 | 0.24 |
| Conditional R<sup>2</sup> | 0.34 | 0.41 |
| . p<0.1; *p<0.05; **p<0.01; ***p<0.001 |  |  |

Finally, also in the model predicting mean VeDBA the interaction term between harness type and species was not significant [*χ*^2^ = 9.06, *p* = 0.06]. Overall, all birds showed a statistically significant decrease in VeDBA (−8%) when equipped with a leg-loop compared to a backpack [leg-loop = −0.08 ± 0.22] (Table 5).

**Table 5:** Output of the LMM with the log of the mean VeDBA included as dependent variable, number of fixes, hour of the day, harness type and species as fixed terms, individual identity and date as random intercepts. The model included only passive flight and was run on a subset of the dataset (every second observation was discarded) to reduce temporal auto-correlation. The interaction term between harness type and species was only slightly significant and therefore excluded.

|  | Passive segments |
| --- | --- |
| **Fixed effects** |  |
| *Estimate (St. Err.)* |  |
| Intercept | -1.81 (0.12)*** |
| Leg-loop | -0.08 (0.22)*** |
| Tawny eagle | -0.14 (0.14) |
| Griffon vulture | -0.56 (0.13)* |
| Himalayan vulture | -0.62 (0.13)* |
| Rüppell's vulture | -0.50 (0.15). |
| Hour | -0.03 (0.007)*** |
| Number of fixes | 0.0007 (0.0001)*** |
| **Random effects (N. groups)** |  |
| *Intercept St. Dev.* |  |
| Individuals | 0.10 (10) |
| Date | 0.09 (7) |
| Observations | 1037 |
| Marginal R<sup>2</sup> | 0.29 |
| Conditional R<sup>2</sup> | 0.42 |
| . p<0.1; *p<0.05; **p<0.01; ***p<0.001 |  |

In three of the seven models (vertical speed and height a.s.l. during soaring and horizontal speed during gliding), the effect size associated to the harness type was higher than the among-individuals and among-dates variability (intercept standard deviation) (Tables 1,2,4). This suggests that the statistically significant variance which we found in at least some of the flight parameters, associated with the harness type, is higher than the variance encountered between individuals and could therefore be relevant from a biological perspective.

### Analysis of the flight sessions

The unit of this analysis was the flight session, therefore it was only applied to the treatment group, as the control individual was only tracked for two flight sessions. The dataset contained a total of 92 observations, where each observation corresponded to one flight session, whose performance were summarised in terms of: total flight duration, total distance covered during the flight, proportion of soaring flight along the track, proportion of active flight and cumulative VeDBA. We applied onesided Wilcoxon test (greater) and found that the difference in flight parameters between harness types was never significantly higher than the baseline, except in the case of the proportion of active flight. In this case, the difference in the proportion of active flight performed with one or the other harness type was significantly higher than the baseline [one-sided Wilcoxon test: V = 40456, *p* = 0.0002]; the mean of the difference between groups was positive, meaning that birds wearing backpacks spent a higher proportion of time using active flight compared to birds wearing leg-loops.

## Discussion

In this study we compared the effect of leg-loop and backpack harnesses on the flight performance of 10 individuals from five raptor species, in a unique setting that allowed us to minimize confounding factors related to environmental context, individual behaviour and handling stress. To our knowledge, this is the first cross-species comparison of the effect of two harness types on fine-scale flight performance. During the analysis we accounted for the animal’s flight behaviour, and analyzed flight performance at the scale of the behavioural segments as well as at the scale of the flight session.

At the level of the behavioural segment, the control individual showed no difference in the flight parameters collected simultaneously by the two harness types, showing that the information we collected were not likely to be affected by the positioning of the device on the animal’s back. The results of the models investigating the effect of harness type on the treatment individuals showed differences in flight performance associated to the two harness types, that suggest a lower drag associated with leg-loop compared to backpack harnesses. In particular, our models showed that birds equipped with leg-loops climbed up to 0.65 ms^−1^ faster and reached heights 19% higher while soaring. A decreased drag associated with the use of leg-loops was also suggested by a lower rate of sinking while gliding and a slightly higher glide ratio, both suggesting that birds equipped with leg-loops could cover a higher horizontal distance per unit of drop in height. Birds wearing leg-loops also showed a higher horizontal speed while soaring and a lower VeDBA, which suggests a lower energy expenditure. However, the variability of these last four parameters associated to the use of leg-loops was comparable to the inter-individual variability; therefore the observed difference in these parameters between the two harness types might not be biologically relevant. Most species equipped with backpack, except for the black kite, showed a higher horizontal speed while gliding, although only for two species this difference was significant. The higher horizontal speed was associated with, and probably offset by, a higher sinking rate, which is probably why backpacked individuals resulted in a similar or slightly lower glide ratio compared to birds equipped with leg-loops. Differences in horizontal speed might also result from different wind conditions, which were not measured. Although we did not have access to high resolution wind information to compare airspeed between harness types, we included date as random intercept in the models as an attempt to control for differences in atmospheric conditions at least between the days. At the level of the flight session, birds wearing leg-loops seemed to spend less time using active flight compared to individuals wearing backpacks, but no other differences were detectable in any of the other flight parameters. A lower proportion of active flight should correspond to a lower energy expenditure during the flight session, although we did not find any difference in cumulative VeDBA between harness types.

Overall, most of our results showed lower flight performance associated with the use of backpack harnesses, probably as a consequence of additional drag caused by the device in its position. This is consistent with a study that visualised the flow over a model penguin, which demonstrated that device-induced turbulence was lower when loggers were placed further back on the body, specifically after the point with maximum girth, where the boundary layer becomes turbulent Bannasch et al. (1994). In our study, the reduction in drag associated with the leg-loop harness resulted in a substantial improvement in flight performance compared to birds with backpacks. For instance, the increase in vertical speed for griffon vultures equipped with leg-loops (0.51 ms^−1^) was 45% of the average vertical speed reported for this species soaring in Israel (1.1 ms^−1^ Harel & Nathan (2018)). It is clear that this could make a substantial difference to the overall cross country speed of these birds given the time they spend in soaring flight (birds in Israel undertook 22.8 thermal soaring cycles per day Harel & Nathan (2018)), even before the improvements in horizontal speed and glide ratio are factored in. We note that other considerations may also affect the optimal logger location, as attaching loggers lower down the back can change the centre of gravity Vandenabeele et al. (2014). This is less likely to be an issue for large birds, such as those in this study, where loggers constitutes a small fraction of their body mass.

The fact that few centimetres difference in the position of the device on the animal’s back could decrease drag with a reduced impact on the birds’ flight performance should encourage the research community to invest more in studying the effect of device attachments. In the last 25 years, several studies highlighted side effects of backpack harnesses on terrestrial bird species (Booms et al. 2011, Stahlecker et al. 2015, Paton et al. 1991, Steenhof et al. 2006, Naef-Daenzer 2007). Our results add to the existing literature in support of considering leg-loops as a good alternative to backpack harnesses, at least for the raptor species investigated in this study. In addition to the positive effect on the birds’ flight performance, suggested by our results, the design of leg-loops has other clear advantages. Leg-loops leave wings, flight muscles and major fat deposits untouched (Naef-Daenzer 2007, Thaxter et al. 2014) and they reduce the risk of entanglement as, in case of damage, they fall off. Leg-loop harnesses are also faster to fit on birds, reducing handling time (especially important when handling wild species), and potentially their stress level. Finally, leg-loops require less material, hence reducing the overall weight of the harness.

Our results suggested no apparent detrimental effect of leg-loop harnesses, but the data used in this study are based on a limited period of data collection and captive individuals. We therefore did not investigate other important parameters such as change in the individual’s behaviour before and after equipping the animals with harnesses, nor potential long-term effects on the individuals’ reproductive success and survival. These potential effects have to be investigated independently, as they cannot be excluded based on results related to flight parameters only. The experience gained with long-term studies using a specific harness type is also useful to evaluate technical improvement. One study, using leg-loops on seabirds, reported that due to the tag position on the animal’s back, the solar panel was covered by feathers and could not charge the device’s battery (Thaxter et al. 2014). In our study we used devices without solar panels, and we could therefore not investigate such technical problems. However, we are aware of long-term tracking studies on griffon vultures using solar-powered tags fitted as leg-loops (Fluhr et al. 2021, Monsarrat et al. 2013, Phipps et al. 2019), as well as a few other ongoing studies with large soaring raptors wearing leg-loop mounted GPS devices. We thus think that technical problems related to energy harvesting can be species specific and in many cases overcome, maybe even reduced through the mere use of leg-loops, at least within the limits posed by the local atmospheric conditions (e.g. hours of sun) and the species-specific behaviour (e.g. time spent flying) and plumage.

Investigating the effect of harness type on fine-scale flight parameters is also relevant in the context of data standardization and comparability (Curk, T., Scacco, M. et al. 2021). The measures of flight performance investigated in our study are commonly used parameters in movement ecology studies focusing on comparing flight behaviour and performance across species, populations or environmental contexts. The data used in such studies are often collected by different research groups using different devices with possibly different attachment methods. It is therefore of primary importance to investigate how the methodology used to measure these information affects the collected data. Not only to the benefit of the animals’ welfare, but also to avoid systematic bias in our results, which would invalidate data comparability and lead to misinterpreting the behaviour we are trying to measure (Curk, T., Scacco, M. et al. 2021, Barron et al. 2010).

## Conclusions

Bio-logging devices are indispensable in movement ecology research, but comparative studies investigating the effect of different device attachments are rare. The available harness types differ in terms of the body parts they restrict, in how easily they can move or fall off and in the resulting position of the device on the animal body, which can in turn affect the device’s drag. The results of our study showed that in large terrestrial species, leg-loop harnesses can be advantageous not only in terms of their design but also because of the reduced drag imposed to the birds, which results in better fine-scale flight performance, and are therefore a good alternative to the commonly used backpack harnesses.

The awareness and quantification of the bias caused by different attachment types will not only benefit our study species, but also allow our research community to make best use of existing data and gain better and more complete insight into the movement ecology field, by using larger sets of data and taking advantage of the comparative aspect that meta-analyses can provide.

## Methods

### Data collection

The work was conducted in Rocamadour, France at the *Le Rocher des Aigles* falconry centre (44.801962°N, 1.612855°E). This study site overhangs a 120 m-deep canyon, providing natural soaring conditions for raptors. Each animal, trained with falconry techniques for the public shows, was released from their perch and flew freely three times a day (at 10:00, 12:00 and 14:00, local time). After their release, the birds usually took-off immediately and had the possibility to fly for about 1 hour (with an average flight duration of 41 minutes) to a maximum distance of 12.8 Km from the releasing point [764.9 m ± 29.4 (mean ± st.err.)]. Between the 28th of June and the 1st of July 2018, we collected GPS and ACC data on 10 individuals from five raptors species: Eurasian griffon vulture (n=4), Rüppell’s vulture (n=1), Himalayan griffon vulture (n=2), tawny eagle (n=2) and black kite (n=1). During each flight, we recorded the time of departure and return of each individual to later isolate only GPS and ACC data collected during the flight sessions.

### Devices and harness types

The devices (70 g weight) were fastened with Velcro on a small aluminium plate and attached to the birds’ body using a Teflon-nylon harness. The total weight of transmitter, aluminium plate and harness was 90 g. The harness was fitted to the birds either as a leg-loop or as a backpack. Backpack harnesses were looped around the bird’s wings with the two loops crossing on the sternum, and the device positioned on the animal’s back between the scapulas (thoracic X-strap harness, described by Bildstein, Botha and Lambertucci (Anderson et al. 2020)). Leg-loop harnesses were looped around the bird’s thighs and the device positioned on the animal’s lower back, on the pelvis above the tail (Anderson et al. 2020).

We used GPS-ACC devices (Technosmart, IT) of different generations. Some devices had GPS and accelerometer sensors separated into two units: *Gipsy 1* (n=8) and *Gipsy 5* (n=1) recorded GPS locations at 4 Hz, and were associated with either *AXY 1* (n=4) or *AGM* (n=3) sensors, which collected ACC data at 25 Hz. Finally *Axytreck* devices (n=3) collected both 1 Hz GPS and 25 Hz ACC. All devices recorded GPS and ACC information continuously. At the beginning of each day, all tags were positioned on a wooden slat to be switched on and calibrated simultaneously.

### Control group

We used as control group data collected from one Eurasian griffon vulture during one day. During that day and two flight sessions, this control individual was equipped simultaneously with both backpack and leg-loop. Both devices measured the same behaviour at the exact same time, and the GPS and ACC devices deployed were of the same generation (*Gipsy 1* and *AXY 1*). Therefore, we expect that potential differences between the flight parameters measured using the two harness types should be purely methodological and associated to the position of the device on the animal’s body. This allowed us to assess if, for the same given behaviour, the position of the device on the animal’s back could affect the information we collect.

### Treatment group

We randomized the combination of device and harness type associated to each individual, to disentangle potential effects associated to the device type, the harness type and the individual behaviour. Each individual bird could thus experience both types of attachment and different devices. Thus, each flight session of the day was considered as a separate unit and during each flight session, individuals were equipped with either a leg-loop or a backpack harness.

### Data processing and behavioural segmentation

The original dataset included 10 individuals from five species and a total of 96 flight sessions (40 with backpacks and 56 with leg-loops). Within each flight session, ACC and GPS data were recorded continuously. ACC data were collected at 25 Hz; GPS data at 1 and 4 Hz depending on the device generation, but they were all sub-sampled to 1 Hz (one GPS fix per second).

We used ACC data to identify active flight. We first calculated the static component of acceleration by taking running means (smoothed values) of the raw acceleration values of each of the three axis over a period of 0.5 seconds, corresponding to two complete flapping cycles (we observed an average of four flapping cycles per second) (Shepard et al. 2008). We then obtained the dynamic component of acceleration by subtracting the smoothed values from the raw values. We finally used the dynamic acceleration of the three axes to derive the VeDBA (Williams et al. 2015, Wilson et al. 2020). We averaged the VeDBA values per second and applied a K-means clustering algorithm with k=2 to distinguish between active and passive flight. Average VeDBA values and activity classes were then associated to the GPS location matching in time.

To segment the GPS data, we applied a running mean of 15 s on the vertical speed; we then applied K-means clustering with k=2 on the smoothed vertical speed to distinguish soaring from gliding behaviour. Vertical speed, horizontal speed and step length between consecutive GPS fixes were calculated for each flight session separately using the R package `move` (Kranstauber et al. 2020).

The results of the two K-means clusterings, the one based on the smoothed VeDBA and the one based on the smoothed vertical speed, were finally combined in one variable with four classes: passive soaring, passive gliding, active soaring and active gliding. The results of the segmentation procedure were inspected visually by plotting the raw ACC values of the three axes and the GPS trajectories in three dimensions.

### Datasets

We analysed the effect of harness type on the flight parameters measured at two different levels. We first focused on the level of the behavioural segment: consecutive GPS fixes belonging to the same behavioural class were assigned to the same segment ID, and their flight parameters averaged across the segment. Therefore, each entry of the dataset used in the analysis corresponded to one behavioural segment with the following associated parameters: mean vertical speed, mean horizontal speed, glide ratio (ratio between the distance covered in the horizontal plane and the distance dropped in height during each gliding segment), maximum height above sea level (a.s.l.) and mean VeDBA. The segments were highly variable in terms of their duration (number of consecutive fixes). To improve comparability of the flight parameters across segments of different duration we excluded segments longer than 733 fixes (>0.01% percentile). This dataset included both the control (1 individual) and the treatment groups.

We then worked at the level of the flight session. Each observation of this dataset corresponded to one flight session, whose performance was summarised in terms of: total flight duration, total distance covered during the flight, proportion of soaring flight along the track, proportion of active flight and cumulative VeDBA. The control individual was excluded from this dataset, as it was only tracked for two flight sessions.

### Analysis of the behavioural segments

The average horizontal speed associated to the segments included in the analysis had a bi-modal distribution, with medians at 0.35 ms^−1^ and 11.40 ms^−1^, and a clear natural divide at 4 ms^−1^. We thus used a 4 ms^−1^ threshold to separate low from high speed segments [max. speed in low speed segments: 3.28 ms^−1^; min. speed in high speed segments: 4.59 ms^−1^]. The segments associated to very low speeds occurred during flight and could not be associated to a specific behaviour. For the following analysis we therefore considered only high speed segments (with average horizontal speed > 4 ms^−1^).

Control and treatment groups were analysed separately.

For the control individual, we used two-sided Wilcoxon signed rank tests to assess if the differences in mean vertical speed, mean horizontal speed, glide ratio, maximum height a.s.l. and mean VeDBA measured using the two harness types was significantly different from 0.

For the treatment group, we used linear mixed-effects models (LMM) (R package `lme4`) (Bates et al. 2015) to test the effect of harness type on the flight performance parameters measured at the level of the flight segments. Mean vertical speed, mean horizontal speed, glide ratio, maximum height a.s.l. and mean VeDBA were used as response variables. As vertical speed, horizontal speed and height a.s.l. are known to differ between the soaring and gliding phases, we tested each of these three flight parameters separately, once during soaring and once during gliding. In contrast, as both soaring and gliding phases are expected to result in a similarly low activity level of VeDBA, we ran only one model for all passive flight segments testing for differences in VeDBA in attachment types. Glide ratio was only analysed for gliding segments. We found unrealistically high glide ratios (between 100 and 914)) to be associated with very low sinking rate (mean vertical speed > −0.16 ms^−1^, more similar to horizontal flight than gliding); we therefore included in the glide ratio model only gliding segments with vertical speed < −0.2 ms^−1^.

In all models, harness type and species were included as interacting categorical predictors, to account for potential differences in the way the different species were affected by the two harness types. Using ANOVA, we assessed the statistical significance of the interaction term and of the harness type, by comparing the full model with null models not including these terms. Hour of the day (with 0 centered at 12:00 UTC) was also included as predictor in all models to acknowledge changes in flight parameters at different times of the day. Finally, we included the segment length (number of fixes in the segment) to account for the variability in the duration of the behavioural segments. Date of the flight session and individual identity were included as random terms in all models.

The height a.s.l. and VeDBA models were run on a subset of the dataset, including one every second observation to reduce temporal auto-correlation. The variable horizontal speed during soaring was square-root transformed while the variables height a.s.l. and VeDBA were log transformed and all models were fitted with a Gaussian error distribution.

### Analysis of the flight sessions

We used non-parametric Wilcoxon tests on the treatment individuals to compare the measured flight parameters between harness types. Specifically, for each species *α* and for each flight parameter *P*, we computed the absolute difference between all combinations of observations of backpack (*BP*) and leg-loop (*LL*). This difference was defined as:

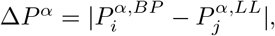

where *i* and *j* represent the *i^th^* and *j^th^* observation (flight session) associated to each harness type. To avoid replicates, we ensured that the number of observations was equal between the two groups: when the number of observations was higher for one of the two harness types, we randomly sub-sampled the number of observations associated to the second harness type.

We then tested whether the distribution of absolute differences between the groups (Δ*P^α^*) was higher (one-sided Wilcoxon test) than the mean of absolute differences within groups (baseline). The baseline *B* was defined as:

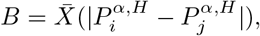

where *H* represents the respective harness type and *α* the species, as the baseline was calculated within species and within harness type.

Data processing and analysis were performed in R (R Core Team 2020).

### Ethic statement

The study was conducted under the permit for equipping vultures with loggers as part of the licence granted to O. Duriez from the Research Centre for Bird Population Studies (CRBPO) of the Natural History Museum (MNHN, Paris). According to the French law of 22 September 2008, the CRBPO has the delegation by the Ministry of Ecology, Energy, Sustainable Development and Land Settlement for allowing the owners of a general bird ringing licence to capture and handle birds from protected species and mark them (with rings or any devices like loggers). The study was conducted under a formal agreement between the animal rearing facility (Rocher des Aigles) and CNRS. Birds were handled by their usual trainer, under the permit of the Rocher des Aigles (national certificate to maintain birds “Certificat de capacité” delivered to the director, Raphaél Arnaud on 4 November 1982). Care was taken to minimize discomfort to the birds and loggers were removed promptly after flights.

## Availability of data and materials

The data that support the findings of this study and the R scripts used to process and analyse the data are available at https://doi.org/10.5281/zenodo.5531226.

## Authors’ contributions

MS, KS and OD designed the study. MS and OD collected the data. AL and MS analysed and interpreted the data. AL and MS wrote the first draft of the manuscript. KS, ES, OD and MW provided valuable comments on the manuscript. All authors read and approved the final version of the manuscript.

## Acknowledgements

We would like to thank the falconry centre *Le Rocher des Aigles*, particularly directors R. Arnaud and D. Maylin, for their welcome and financial support, and all bird trainers for their technical help. We also acknowledge Camille Nouis for her help during the data collection and the preprocessing of the data. We are also thankful to Daniel Hegglin for the discussions and advice on leg-loop attachment in large soaring birds.

